# Isofagomine inhibits multiple TcdB variants and protects mice from *Clostridioides difficile* induced mortality

**DOI:** 10.1101/2023.09.19.558375

**Authors:** Ashleigh S. Paparella, Isabella Brew, Huynh A. Hong, William Ferriera, Simon Cutting, Farah Lamiable-Oulaidi, Michael Popadynec, Peter C. Tyler, Vern L. Schramm

## Abstract

*Clostridioides difficile* causes life-threatening diarrhea and is the leading cause of healthcare associated bacterial infections in the United States. During infection, *C. difficile* releases the gut-damaging toxins, TcdA and TcdB, the primary determinants of disease pathogenesis and are therefore therapeutic targets. TcdA and TcdB contain a glycosyltransferase domain that uses UDP-glucose to glycosylate host Rho GTPases, causing cytoskeletal changes that result in a loss of intestinal integrity. Isofagomine inhibits TcdA and TcdB as a mimic of the oxocarbenium ion transition state of the glycosyltransferase reaction. However, sequence variants of TcdA and TcdB across the clades of infective *C. difficile* continue to be identified and therefore, evaluation of isofagomine inhibition against multiple toxin variants are required. Here we show that Isofagomine inhibits the glycosyltransferase activity of multiple TcdB variants and also protects TcdB toxin-induced cell rounding of the most common full-length toxin variants. Further, isofagomine protects against *C. difficile* induced mortality in two murine models of *C. difficile* infection. Isofagomine treatment of mouse *C. difficile* infection permitted recovery of the gastrointestinal microbiota, an important barrier to prevent recurring *C. difficile* infection. The broad specificity of isofagomine supports its potential as a prophylactic to protect against *C. difficile* induced morbidity and mortality.

## Introduction

*Clostridioides difficile* is a Gram-positive spore forming bacterial pathogen that causes pseudomembranous colitis and is the leading cause of healthcare associated infections in the United States^1,2^. *C. difficile* infections (CDI) typically occur following treatment with broad-spectrum antibiotics that cause dysbiosis of the normal gastrointestinal (GI) microbiota, allowing *C. difficile* spores to geminate and proliferate in the colon. *C. difficile* releases up to 3 toxins that result in inflammation of the colon and cause life-threatening diarrhea^3,4^. Current treatment for CDI involves administering broad-spectrum antibiotics including vancomycin or fidaxomicin. However, antibiotic therapy prevents re-establishment of a healthy GI microbiota and therefore relapse of CDI occurs in approximately 20% of patients^5-7^. Furthermore, *C. difficile* has acquired resistance to multiple antibiotic classes, highlighting the urgency for new treatment options^8^. One emerging strategy to treat CDI is to develop therapeutics that target the gut-damaging toxins secreted by *C. difficile* during infection. Targeting the toxins directly minimizes damage to the human colon and spares the human GI microbiota, maintaining a critical barrier to CDI^7^. The major toxins secreted by *C. difficile* during infection are TcdA (308 kDa) and TcdB (270 kDa). These multi-domain protein toxins are the primary determinants of disease pathogenesis^9,10^. A third toxin, CDT binary toxin is produced by a minority of *C. difficile* strains during infection and can also contribute to disease when present^11,12^. The mechanism of action for TcdA and TcdB has been reviewed extensively^13-16^. TcdA and TcdB first bind to target cells via a C-terminal receptor binding domain that triggers internalization by clathrin-mediated endocytosis. Next, acidification of endosomes causes pH-dependent conformational changes leading to the formation of a membrane pore and subsequent delivery of the N-terminal auto-processing cysteine protease domain (CPD) and glycosyltransferase domain (GTD) into the cytosol. The CPD is allosterically activated by intracellular inositol hexakisphosphate, catalyzing the release of the GTD into the cytosol. There, the GTD glycosylates and inactivates Rho GTPases, including Rac1 and Cdc42 at Thr35 and RhoA at Thr37 in the switch I effector region, using UDP-glucose as a glycosyl donor (Figure 1). TcdA/B glycosylation of Rho GTPases leads to actin-depolymerization resulting in a loss of structural integrity of the cell and activation of caspase-3 and caspase-9 dependent pathways, followed by cell death^13-16^.

**Figure 1:**
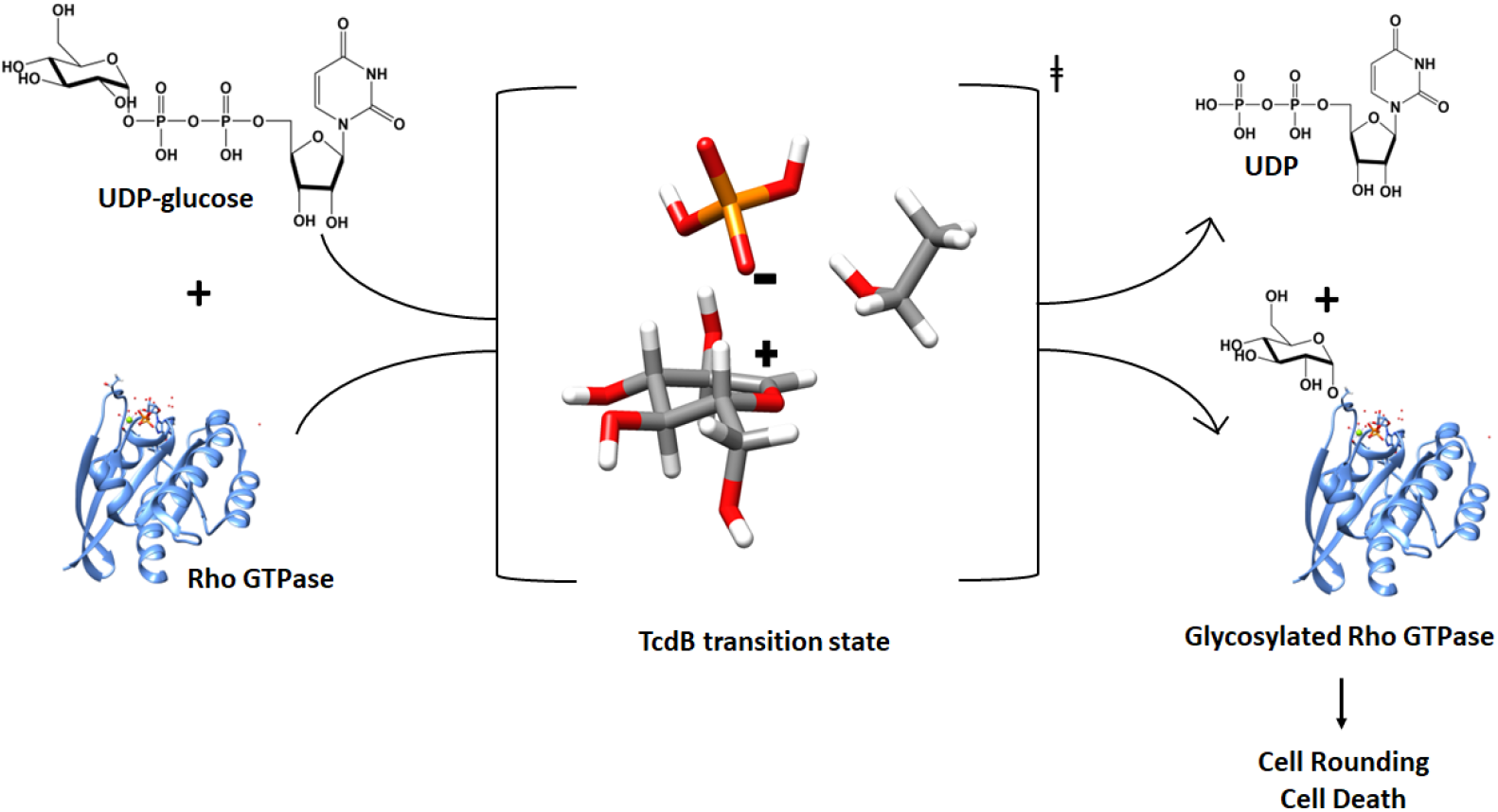
Schematic of TcdA and TcdB glycosyltransferase reaction. (Adapted from Ref 25)

*C. difficile* disease pathogenesis is complicated by diverse *C. difficile* strains found to express TcdA and TcdB with sequence variations from the reference strain VPI10463 (designated TcdA1 and TcdB1)^17^. Two recent studies have proposed grouping TcdB toxin variants into either 8 or 12 subtypes (TcdB1-B12)^18,19^. Compared to TcdB, TcdA showed far fewer variations in both studies^17,18^. Sequence variations in the toxin genes could influence receptor binding, specificity towards Rho GTPases, antigenicity and overall toxicity. Furthermore, TcdB variants have been shown to use different receptors to gain entry into mammalian host cells, including the CSPG4, Frizzled and TFPI protein receptors^20,21^. Given that multiple TcdA and TcdB variants have been identified, it is essential to evaluate TcdA and TcdB targeted therapeutics for inhibition of multiple variants.

One recently approved *C. difficile* therapeutic is Bezlatoxumab (Zinplava™), a monoclonal antibody that acts by recognizing two epitopes in the C-terminal receptor binding domain of TcdB1^22^. Given that TcdB variants use different host cell receptors, antibody therapeutics designed to neutralize the TcdB toxins via binding to the receptor binding domain, may not recognize all TcdB variants. Indeed, it has been shown that antibodies directed against the C-terminal receptor binding domains show different neutralizing activities to TcdB1 and TcdB2^23^. Therefore, developing therapeutics that inhibit all TcdB variants with similar levels of potency is important for the development of new TcdA and TcdB therapeutics.

Previously, we reported isofagomine as an inhibitor of the GTD’s of TcdA1 and TcdB1^24^. Isofagomine is a transition state analogue that mimics the positively charged glucocation formed at the transition state of the TcdA and TcdB glycosyltransferase reactions^25^. Isofagomine binds in the TcdA and TcdB active sites and forms an ion-pair interaction with the negatively charged β-phosphate of UDP, a by-product of the Tcd glycosyltransferase reaction (Figure 2)^24,25^. To further characterize isofagomine as a new *C. difficile* therapeutic, we report the inhibitory activity of isofagomine against 8 variants (TcdB1-8) of the TcdB-GTD. We also compared the efficacy of isofagomine to prevent mammalian cell rounding induced by the 4 most common full-length variants of TcdB (TcdB1-4). Finally, we demonstrate that isofagomine can protect against *C. difficile* induced mortality and allows the recovery of the GI microbiota in a mouse model of CDI.

**Figure 2:**
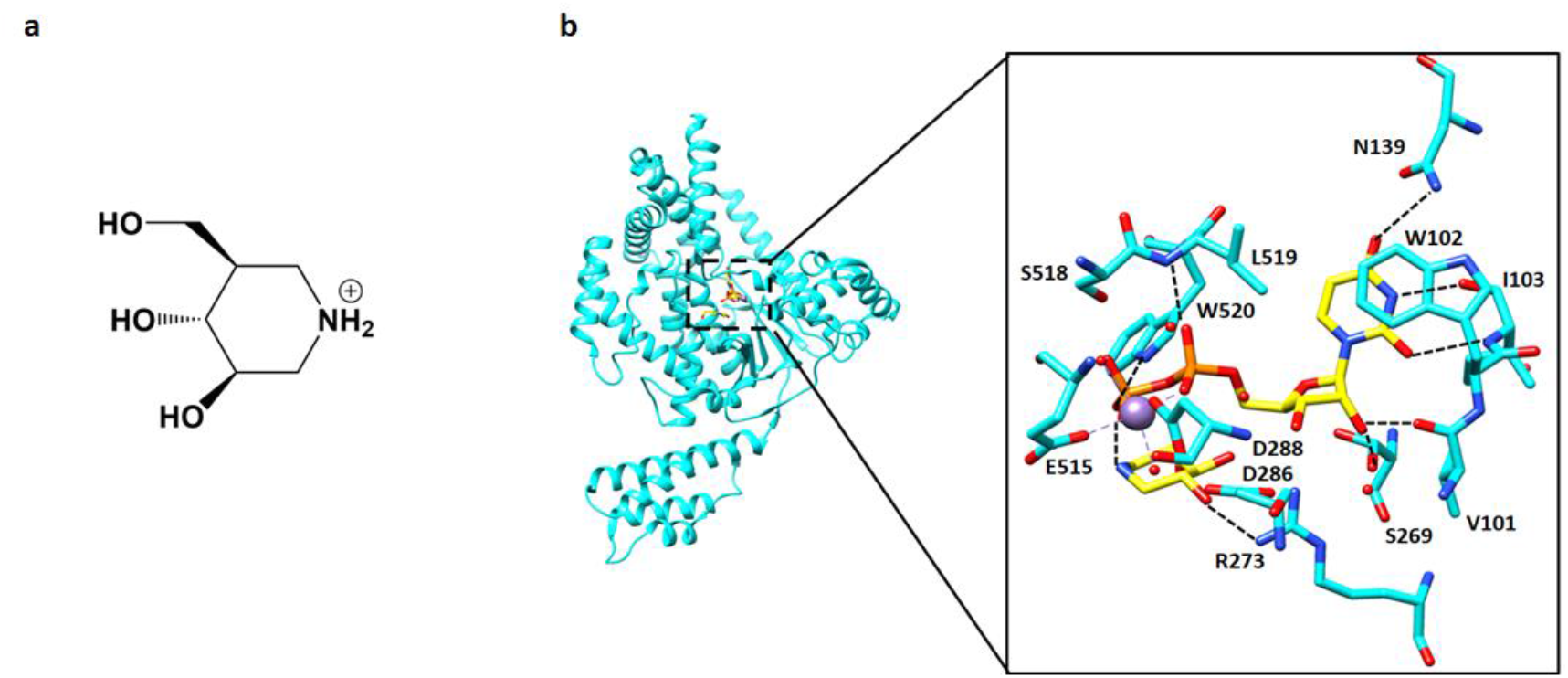
Chemical structure of isofagomine and summary of interactions in TcdB1 active site: **a)** Chemical structure of isofagomine. **b)** X-Ray crystal structure of TcdB1 glycosyltransferase domain in complex with UDP and isofagomine (PDB: 7LOU). UDP and isofagomine are colored in yellow and TcdB1 amino acid residues are in cyan. Mn^2+^ metal co-factor is indicated by purple sphere and water molecules are indicated as red dots. Hydrogen bonding interactions are indicated by dashed black lines.

## Results

### Isofagomine inhibits multiple TcdB variants

Isofagomine is an uncompetitive inhibitor of TcdA1 and TcdB1, as it binds TcdA1 and TcdB1 only in the presence of UDP, a product of the TcdA and TcdB glycosyltransferase reactions^24^. Here, we test whether isofagomine exhibits similar inhibitory activity against other TcdB variants. TcdB-GTD variants 1-8 were expressed and purified and kinetic constants for substrates UDP-glucose and Rac1 were determined using a radiolabel incorporation assay which measures the amount of glycosylated Rac1^24,25^. The Michaelis-Menten (*K*_*m*_) constant for UDP-glucose was similar for each variant and ranged from 4.3 μM to 9.4 μM (Table 1). The *K*_*m*_ for Rac1 varied from 13.3 μM to 55.4 μM amongst the TcdB variants (Table 1). The TcdB variants bind to Rho GTPases with different affinities and could show different affinities for other Rho GTPases, such as RhoA, Cdc42. Interestingly, TcdB4 exhibited the largest *K*_*m*_ for Rac1 and exhibited the slowest turnover (*k*_*cat*_ = 0.01 ^s-1^), suggesting that Rac1 may not be the preferred Rho GTPase substrate for TcdB4. The product inhibitory activity of UDP against the GTD’s of TcdB1-8 was measured as another index of catalytic differences (Table 2). UDP inhibited all TcdB variants in the μM range and was most potent against TcdB1 (*K*_*i*_ = 7.9 ± 4.1 μM) and least potent against TcdB7 (*K*_*i*_ = 41.0 ± 0.1 μM). The inhibition constant for isofagomine was measured in the presence of excess UDP (2 x *K*_*i*_) (Table 2). In the presence of UDP, isofagomine inhibited all TcdB variants. With the exception of TcdB4, the *K*_*i*_ for isofagomine varied from 18 nM for TcdB5 to 97 nM for TcdB7. The isofagomine *K*_*i*_ for TcdB4 was 13.8 ± 1.4 μM and was 148-fold less-potent compared to TcdB1. Isothermal titration calorimetry was also used to measure a dissociation constant (*K*_*D*_) of 10.9 ± 5.8 μM for isofagomine binding to TcdB4 in the presence of UDP, similar to the kinetic *K*_*i*_ value (Figure S1).

**Table 1:**
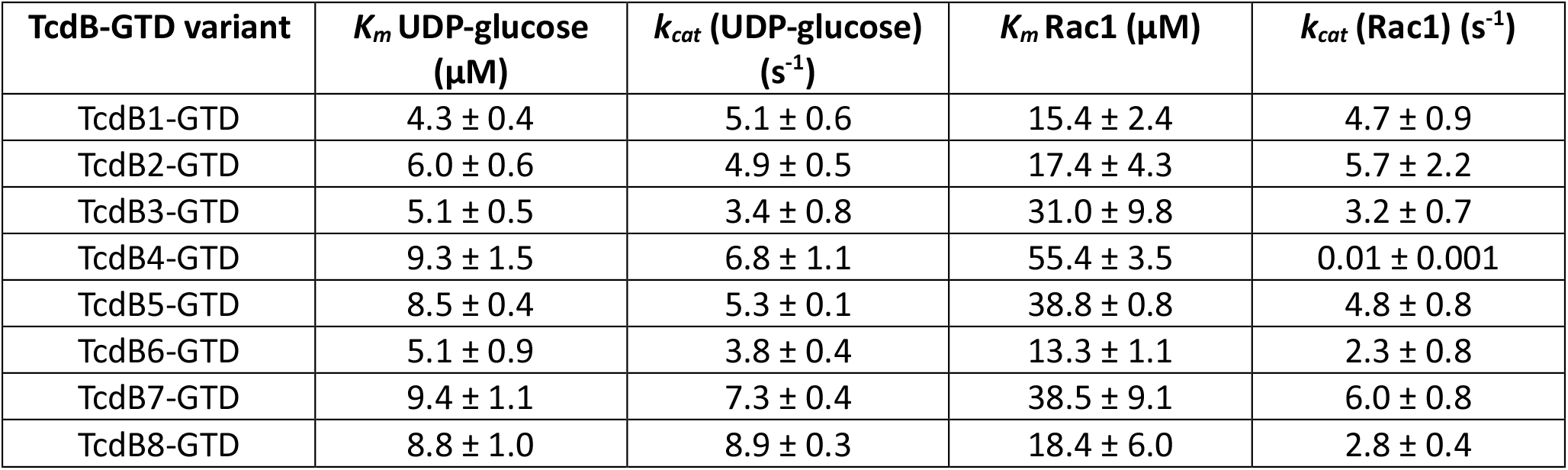
Michaelis-Menten (*K*_*m*_) and Turnover (*k*_*cat*_) numbers for TcdB-GTD 1-8 variants.

**Table 2:**
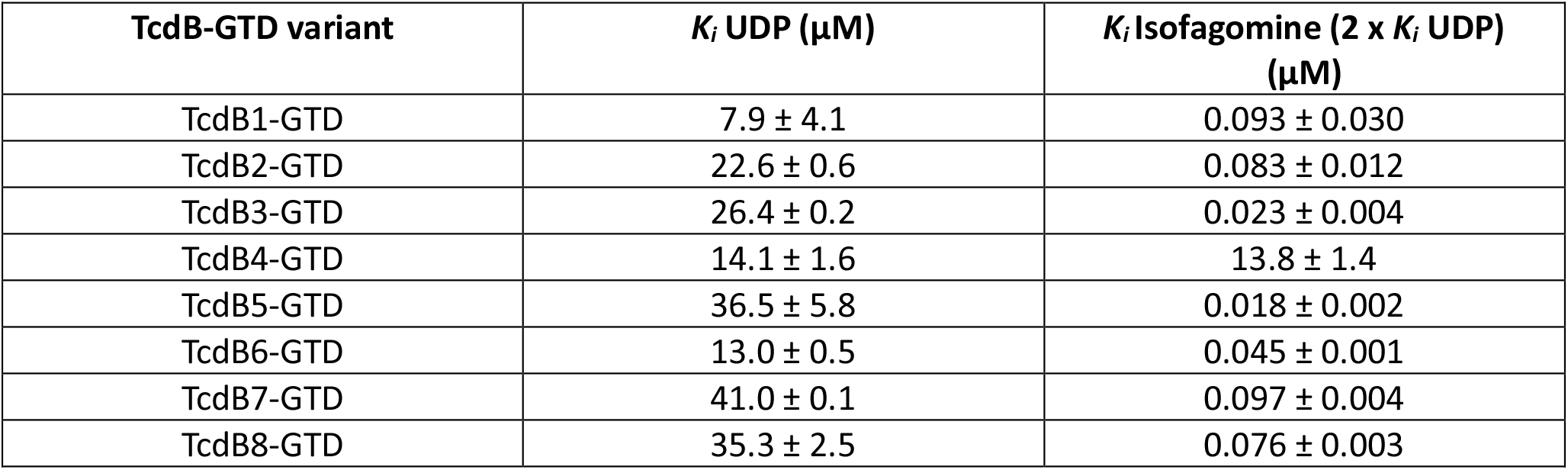
Inhibition data for Isofagomine and UDP against TcdB-GTD glycosyltransferase activity.

| TcdB-GTD variant | $K_i$ UDP ($\mu\text{M}$) | $K_i$ Isofagomine ($2 \times K_i$ UDP) ($\mu\text{M}$) |
| --- | --- | --- |
| TcdB1-GTD | $7.9 \pm 4.1$ | $0.093 \pm 0.030$ |
| TcdB2-GTD | $22.6 \pm 0.6$ | $0.083 \pm 0.012$ |
| TcdB3-GTD | $26.4 \pm 0.2$ | $0.023 \pm 0.004$ |
| TcdB4-GTD | $14.1 \pm 1.6$ | $13.8 \pm 1.4$ |
| TcdB5-GTD | $36.5 \pm 5.8$ | $0.018 \pm 0.002$ |
| TcdB6-GTD | $13.0 \pm 0.5$ | $0.045 \pm 0.001$ |
| TcdB7-GTD | $41.0 \pm 0.1$ | $0.097 \pm 0.004$ |
| TcdB8-GTD | $35.3 \pm 2.5$ | $0.076 \pm 0.003$ |

Finally, we tested the efficacy of isofagomine to protect against TcdB holotoxin-induced HeLa cell rounding. For these assays, we used full-length variants TcdB1-4, these being the four of the most common TcdB variants found in *C. difficile* clinical strains^18^. Isofagomine inhibited TcdB1 induced cell rounding with an IC_50_ of 8.4 ± 1.5 μM, similar to that previously reported for Vero cells^24^. Isofagomine also inhibited TcdB2 and TcdB3 induced cell rounding with IC_50_ values of 17.4 ± 2.9 μM and 8.4 ± 1.5 μM respectively (Table 3). Consistent with the kinetic assays described above, isofagomine also exhibited the weakest inhibitory activity against TcdB4-induced mammalian cell rounding (IC_50_ > 200 μM). Of the four TcdB variants tested, the nucleotide sequences encoding TcdB1 and TcdB2 are the most common in *C. difficile* genomes^18,19^. Overall, Isofagomine inhibits multiple TcdB variants and shows potential to be effective against multiple *C. difficile* strains.

**Table 3:** Efficacy of isofagomine against TcdB induced cell rounding of HeLa cells.

| TcdB variant | Calculated $\text{IC}_{50}$ ($\mu\text{M}$) |
| --- | --- |
| TcdB1 | $8.4 \pm 1.5$ |
| TcdB2 | $17.4 \pm 2.9$ |
| TcdB3 | $8.6 \pm 1.1$ |
| TcdB4 | >200 |

### Isofagomine protects against *C. difficile* induced mortality *in* vivo

Two mouse models were used to investigate if isofagomine protects mice from *C. difficile* induced mortality. The toxigenic model involves an intraperitoneal injection of 2 µg/kg full-length TcdB toxin followed by monitoring for signs of toxicity^26^. Three groups of mice were provided with isofagomine tartrate in the drinking water (20 mg/kg, 60 mg/kg and 180 mg/kg), corresponding to 10 mg/kg, 30 mg/kg and 90 mg/kg isofagomine respectively. Isofagomine treatment commenced 24 h prior to administration of full-length TcdB1 and continued for up to 2 days post challenge (Figure 3a). Mice that received 0, 20, 60 and 180 mg/kg isofagomine tartrate had a 16%, 33%, 50% and 100% survival at 12 hr post toxin administration. At 24 hr, the survival rates were 16%, 33%, 0% and 33% for the same isofagomine doses (Figure 3b). The survival curve for the group that received 180 mg/kg isofagomine tartrate was significantly different to the control group as defined by the Gehan-Breslow-Wilcoxon test which allocates more weight to deaths that occur at earlier time points (p = 0.0350). Although all mice (with the exception of 2 mice at 0 mg/kg and 20 mg/kg isofagomine tartrate) became moribund, treatment with isofagomine tartrate significantly delayed TcdB1 induced toxicity at higher doses.

**Figure 3:**
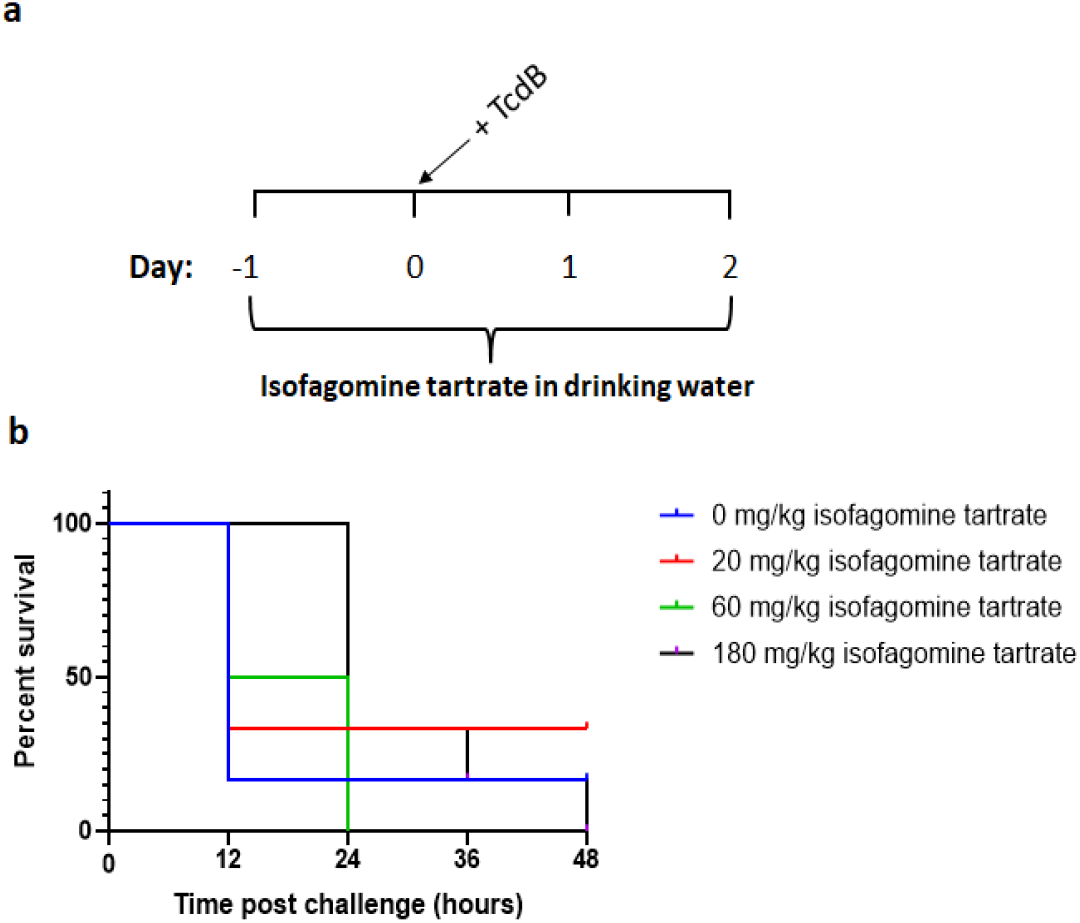
Isofagomine delays TcdB induced toxicity in a toxigenic model of *C. difficile* infection. **a)** Protocol schematic for toxigenic mouse model of *C. difficile* infection. **b)** Survival plots showing the effects of isofagomine treatment (n = 6 mice per group). Values were compiled using a Kaplan-Meier method using animals scored at 48 h as the end point before humane termination.

A second mouse model for isofagomine protection against *C. difficile* toxicity involved bacterial infection of the gut. This mouse model, based on the model described in Chen *et al* more closely mimics the human disease^27^. Here, mice were first exposed to an antibiotic cocktail containing kanamycin, gentamicin, colistin, metronidazole and vancomycin for 3 days to compromise the GI microbiota that is natural barrier to *C. difficile* colonization and infection^7^. After 2 days, mice were treated with clindamycin and administered isofagomine tartrate in the drinking water at 5 mg/kg, 15 mg/kg and 45 mg/kg. These doses correspond to 2.5 mg/kg, 7.5 mg/kg and 22.5 mg/kg of isofagomine respectively. One day later, mice were challenged with *C. difficile* VPI 10463. Isofagomine treatment was continued for 11 days after *C. difficile* challenge (Figure 4a). This treatment regimen provides a protection protocol to block effects of the toxin during active infections. In the Chen *et al* model, mice develop symptoms to reach an infection end point within 2-3 days post-challenge^27^. Typical symptoms include weight loss, hunched posture and diarrhea. In this study, all infected mice, with the exception of the vancomycin control group, lost 10-15% body weight on days 2-6 post-challenge (Figure 4b). Treated mice, with the exception of the 45 mg/kg treatment group, displayed significantly lower diarrhea scores compared to the non-treated mice on day 2 (Figure 4c). Although all isofagomine treated groups exhibited weight-loss and displayed symptoms of CDI, the untreated group reached a clinical end point of 16% survival (1 of 6 mice) on day 2 (Figure 4d). Furthermore, histological analysis of mouse colon tissue indicated that non-treated mice had submucosal edema and loss of epithelial cells lining the mucosal surface (Figure S2). In comparison the histological analysis of vancomycin and 15 mg/kg isofagomine tartrate treated samples indicated normal numbers and appearance of epithelial cells and the submucosa was within normal histological limits (Figure S2). Finally, in the Isofagomine treated groups, 4 mice (67%), 5 mice (83%) and 3 mice (50%) per group survived while receiving 5 mg/kg, 15 mg/kg or 45 mg/kg of isofagomine tartrate respectively (Figure 4e). In the group that received vancomycin, all animals survived since *C. difficile* VPI 10463 is vancomycin-sensitive (Figure 4e). Overall, treatment with 5 mg/kg and 15 mg/kg isofagomine tartrate provided significant protection from *C. difficile* induced mortality.

**Figure 4:**
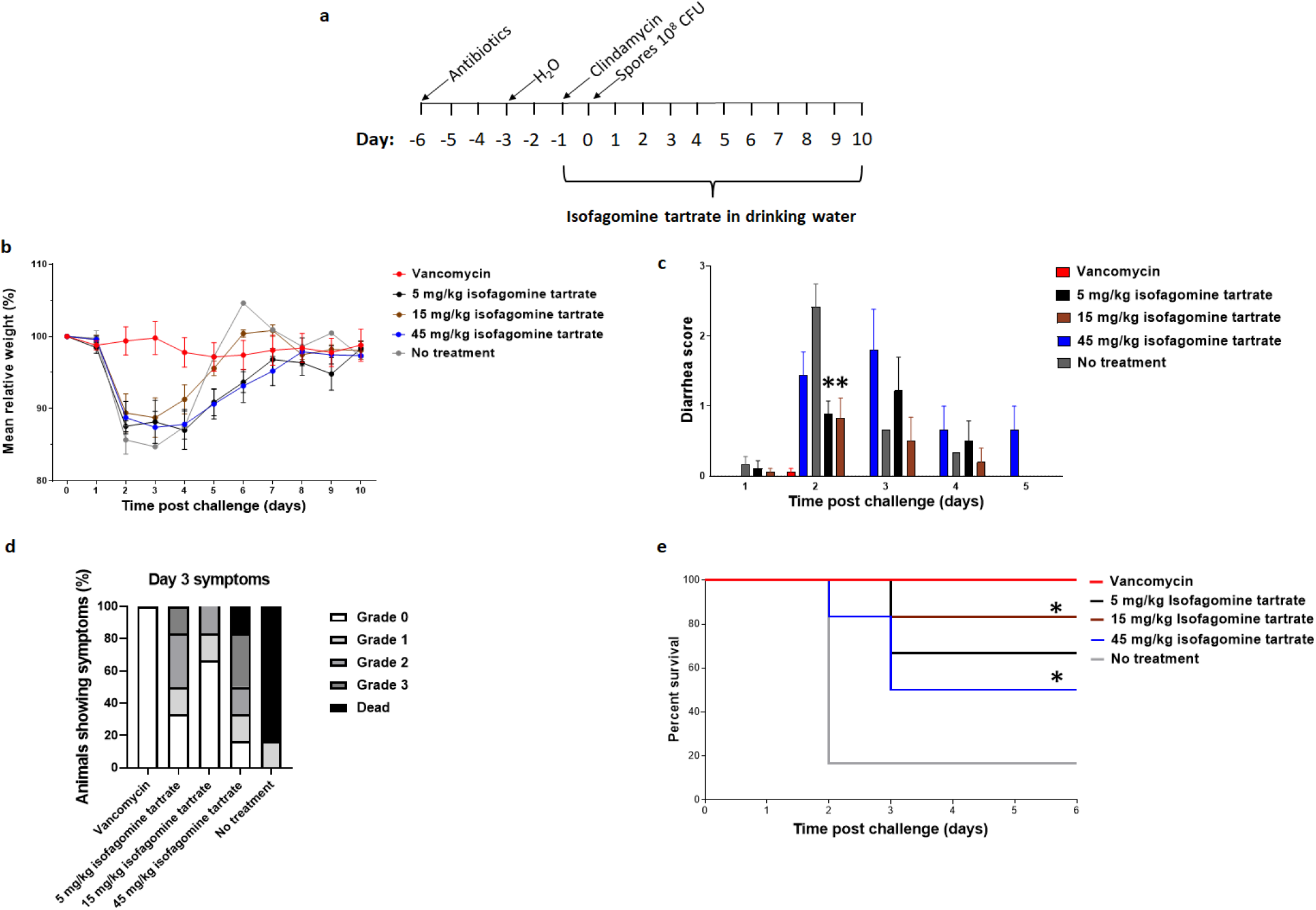
Isofagomine treatment protects mice against *C. difficile* induced mortality. **a)** Protocol schematic for mouse model of CDI. **b)** Weights of mice after challenge with *C. difficile* spores on day 0. Mice were treated with 5 mg/kg, 15 mg/kg or 45 mg/kg isofagomine tartrate. Each data point represents the mean ± SEM. **c)** Diarrhea score of infected mice after 1-day post-challenge, * indicates p < 0.05 as determined by unpaired student t-test. **d)** Symptoms of mice 3 days after challenge with *C. difficile* spores. Grade 0 (hard fecal pellet, normal behaviour), Grade 1 (difficulty in producing feces, mild weakness), Grade 2 (loose stool, weakness) and Grade 3 (liquid feces, inflamed rectum, wet (soiled) tail, extreme weakness). **e)** Survival plots showing the effects of isofagomine treatment (n = 6 mice per group). Values were compiled using a Kaplan-Meier method using animals scored at 6 days post-challenge before humane termination at 10 days post-challenge. * indicates p < 0.05 as determined by Log-rank/Mantel-Cox test.

### Isofagomine treatment allows recovery of gut microbiota

The effects of vancomycin and isofagomine treatment on the GI microbiota during CDI were compared by fecal microbiome analysis. In the Chen *et al* model described above, mice are given an antibiotic cocktail prior to *C. difficile* spore challenge^27^. To compare the microbial diversity pre- and post-antibiotic cocktail, fecal samples were collected at day -6 (pre-antibiotic cocktail) and day -2 (post-antibiotic cocktail) (Figure 5a). Before antibiotic treatment, the microbiota was dominated by S24-7, *Lachnospiraceae, Rikenellaceae* and *Bacteroidaceae* families (Figures 5b-f). The GI microbial diversity was high as determined by the Shannon Index (Figure 5g). Following the administration of antibiotics in the drinking water, the diversity of the GI microbiota significantly reduced as measured by the Shannon index (Figure 5g, p < 0.0001) and the samples were mostly comprised of *Lactobacillaceae* and *Porphyromonadaceae* families (Figures 5b-f). After vancomycin or isofagomine treatment had concluded (day 10), fecal samples were collected from the surviving mice in each group and the GI microbial diversity was also assessed (Figure 5a). The GI microbiota of vancomycin treated mice showed some recovery of microbial diversity (p-value < 0.05), however the microbiota was dominated by *Lactobacillaceae* and *Porphyromonadaceae* families (Figure 5b). The mouse group that received 15 mg/kg isofagomine tartrate exhibited the most robust recovery in microbial diversity and was significantly improved in comparison to vancomycin treatment (p < 0.0001) (Figure 5g). The gut microbiota of mice that received 15 mg/kg isofagomine were dominated by *Lactobacillaceae, Lachnospiraceae, Porphyromonadaceae*, S24-7, *Rikenellaceae* and *Bififobacteriaceae* families (Figure 5e). Overall, the fecal microbiome analysis indicated that isofagomine treatment of CDI permits faster recovery of the GI microbiota than treatment with vancomycin.

**Figure 5:**
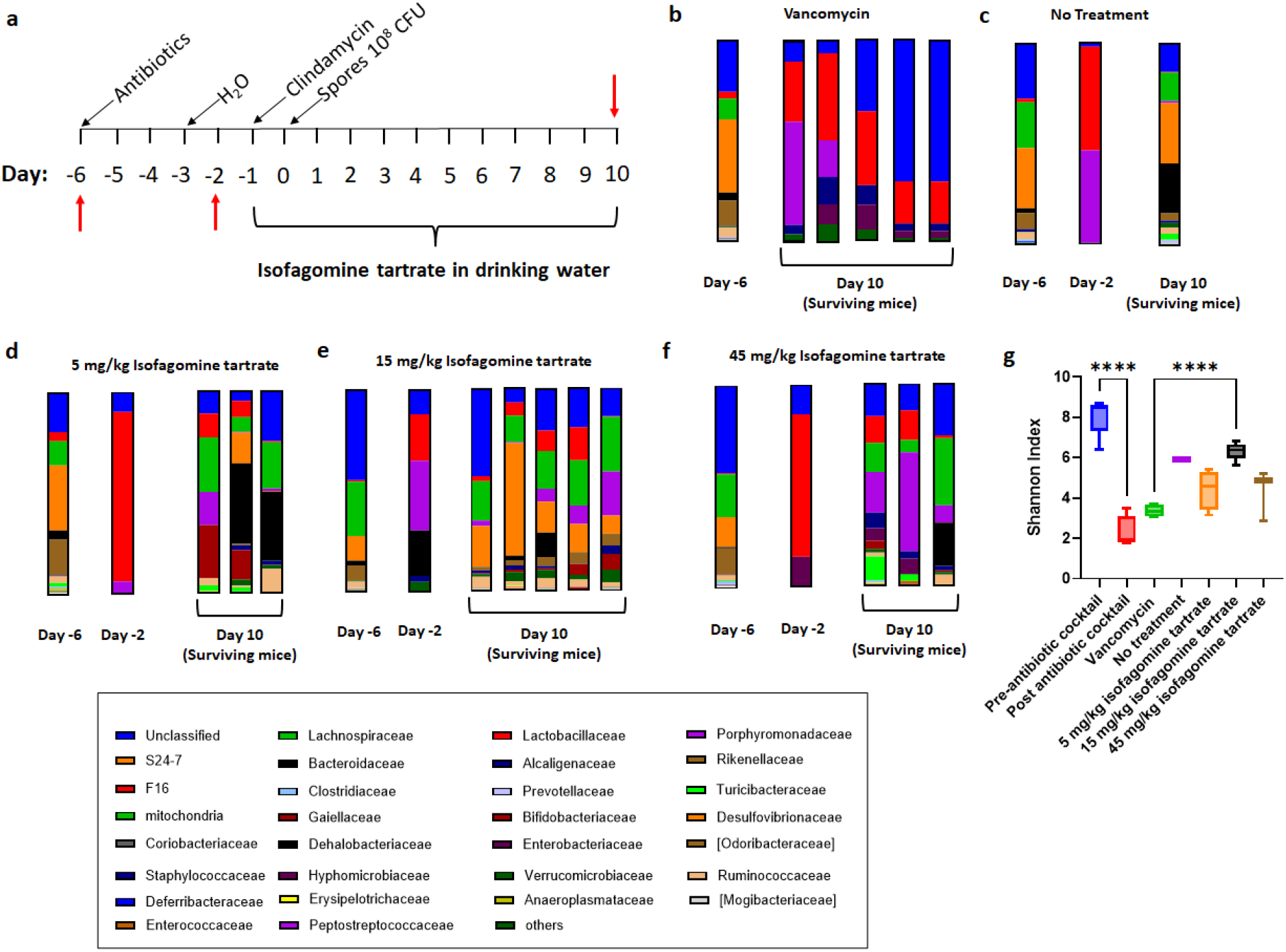
Effect of isofagomine treatment on GI microbial diversity and composition. **a)** Protocol schematic for collection of Fecal sample, red arrows indicate sample collection days. Fecal samples were collected pre-antibiotic cocktail (Day -6), post-antibiotic cocktail (Day -2) and from each surviving animal at the conclusion of isofagomine treatment (Day 10). First sampling day (day -6) is prior to commencement of antibiotic treatment. For figures **b -f** each bar plot represents the mean relative abundance of the top 30 bacterial families identified. **b)** Vancomycin treatment. **c)** Non-treated mice (standard drinking water). **d)** 5 mg/kg isofagomine tartrate. **e)** 15 mg/kg isofagomine tartrate. **f)** 45 mg/kg isofagomine tartrate. **g)** Shannon diversity index. The center line, bounds of box and whiskers of the boxplot represents the mean, inter-quartile and range of sample diversity. p-value was assessed using one-way ANOVA where **** indicates p < 0.0001.

## Discussion

Infections with *C. difficile* typically occur following antibiotic treatment that disrupts the GI microbiota and provides an opportunistic environment for *C. difficile* to populate the colon. Treatment of CDI with further antibiotics does not encourage the re-establishment of the normal GI microbiota and is associated with a high recurrence rate (20%)^7^. Therefore, the development of non-antibiotic therapeutics represents an emerging strategy to combat CDI. TcdA and TcdB are the major virulence factors produced by *C. difficile* during infection and are the primary determinants of disease pathogenesis^9,10^. Therefore, therapeutics that target TcdA and TcdB have the potential to prevent *C. difficile* pathogenesis while allowing the recovery of the normal GI microbiota. However, studies of clinical isolates have identified TcdA and TcdB variants which enter mammalian cells through different receptors and express TcdB of different amino acid sequences. Therefore, small molecule therapeutics that inhibit multiple TcdA and TcdB variants will be needed to use oral small molecule targeting of these toxins.

The efficacy of isofagomine was tested against major and multiple variants of TcdB (TcdB variants 1-8). Isofagomine inhibits the GTD of TcdA and TcdB by mimicking the oxocarbenium ion transition state formed during the TcdA and TcdB GTD reaction^24,25^. Isofagomine inhibited the GTD activity of all TcdB variants tested. However, the relatively rare TcdB4-GTD variant differed substantially from the others, with an inhibition constant over 100-fold greater than the more common TcdB-GTD variants where the inhibition constant was < 0.1 μM. In concert with the low affinity of isofagomine for TcdB4, a decrease in potency of isofagomine against TcdB4 was also demonstrated in mammalian cell rounding assays (IC_50_ > 200 μM). The GTD of TcdB4 shares 78.8% sequence identity to TcdB1-GTD and interestingly shares, 98.9% sequence identity to the GTD of TcdB3, despite there being a 600-fold difference in the inhibition constant for isofagomine. There are 5 amino acid differences between the GTD’s of TcdB3 and TcdB4, the most notable being position 385 within the active site. TcdB3 has an asparagine residue at position 385, TcdB1 has a glutamine residue at position 385, while aspartic acid is in the same position in TcdB4. As Asp385 is near the isofagomine binding site, this substitution is likely responsible for the difference in isofagomine binding affinity (Figure 6). Although isofagomine displayed weaker inhibitory activity against TcdB4, it did exhibit high potency against the two most common TcdB variants TcdB1 and TcdB2^18,19^.

**Figure 6:**
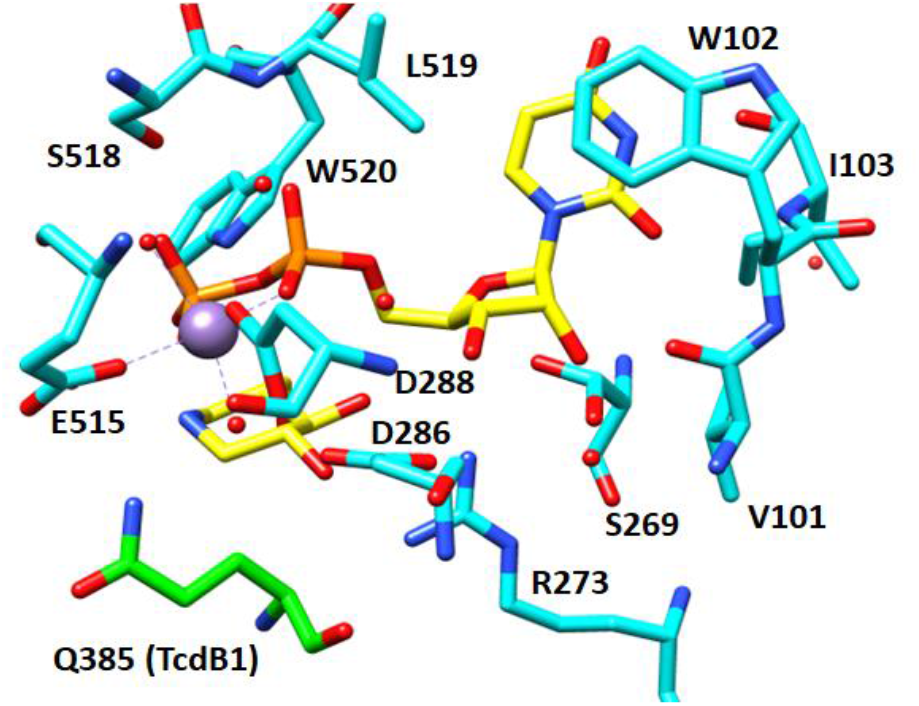
Position of Gln385 in the TcdB1 active site: X-Ray crystal structure of TcdB1 glycosyltransferase domain in complex with UDP and isofagomine (PDB: 7LOU). UDP and isofagomine are colored in yellow and TcdB1 amino acid residues are in cyan. Mn^2+^ metal co-factor is indicated by purple sphere and water molecules are indicated as red dots. Amino acid position 385 (Gln in TcdB1) is indicated in green.

The efficacy of isofagomine was evaluated in two mouse models. In the systemic toxigenic model, mice are given a near-lethal dose of TcdB via intraperitoneal injection. In our study, 83% of mice that did not receive isofagomine tartrate became moribund within 12 hours, while none of the mice that received the highest dose of isofagomine tartrate (180 mg/kg) became moribund within 12 hours. These results indicated that isofagomine tartrate delayed the toxicity of TcdB1. In the *C. difficile* GI infection mouse model, 15 mg/kg of isofagomine tartrate exhibited 83% protection against *C. difficile* induced mortality. Furthermore, treatment of CDI with 15 mg/kg isofagomine tartrate also permitted rapid improvement in GI microbial diversity in comparison to the vancomycin treatment. A healthy GI microbiota is a preventative barrier to CDI and therefore the combination of the inhibition of TcdA/B and recovery of diversity of the gut microbiota highlights isofagomine as a potential anti-*C. difficile* therapeutic. Isofagomine tartrate was previously investigated in human clinical trials as a therapeutic to stabilize the defective glycohydrolases found in Gaucher disease (NCTs: 00433147, 00446550, 00813865, 00875160). Oral doses of isofagomine were well tolerated up to 225 mg/day, however most patients showed no reduction in disease symptoms. Isofagomine trials ceased after phase II clinical trials. Here, we administered isofagomine in the drinking water and therefore mice received continuous doses of isofagomine. Published pharmacokinetic studies in rodents indicate that isofagomine is widely distributed and reaches maximum concentration in the plasma within 1 hour after oral gavage. However, isofagomine is cleared rapidly, with plasma levels decreasing >500-fold from the maximum within 24 hours^30^. Isofagomine prodrugs, slow-release formulations or analogues with longer residence times on the TcdA and TcdB targets should be explored to enhance the clinical potential of isofagomine.

## Materials and Methods

### Gene Expression and Protein Purification

The open reading sequences encoding TcdB1-GTD to TcdB8-GTD were synthesized with a C-terminal His6 tag and cloned into pET28a(+) by Genscript. Sequence length and strain information is shown in Supplementary Table 1. TcdB-GTD variants were expressed in *Escherichia coli* (One Shot BL21 Star (DE3)) cell line. Conditions for protein expression and protein purification of TcdB1-8-GTD were carried out as described previously^24,25^ with purification by Ni-NTA chromatography and protein preparation in 20 mM Tris pH 7.5, 150 mM NaCl, 15% v/v glycerol, freezing and storage at -80 °C.

Expression of Rac1 GTPase (1-192) was carried out in *E. coli* (One Shot BL21 Star (DE3) cell line and also followed the method of Paparella *et al*^24,25^. The gene encoding Rac1 was synthesized with an N-terminal His6 tag and cloned into pD451-SR:391299 plasmid (Atum Bio). Purification of Rac1 was performed by Ni-NTA chromatography and protein preparation in 20 mM Tris pH 7.5, 150 mM NaCl, 15% v/v glycerol, freezing and storage at -80 °C.

The genes encoding full-length TcdB toxin variants (TcdB1,3 and 4) were synthesized with a C-terminal His-tag and cloned into pC-His 1622 by Genscript. The pC-His1622 vector was purchased from MoBiTec. TcdB2 was kindly provided by Jimmy D. Ballard^17^. Expression and purification of full-length TcdB variants was carried out as described previously in Tam *et al*^28^. Briefly, transformed *Bacillus megaterium* was inoculated into LB containing tetracycline (12.5 μg/mL) and grown at 37 °C to an OD_600_ of approximately 0.3 -0.4, followed by overnight xylose induction (0.5%) at 37 °C for TcdB1 and 25 °C for TcdB3/4. Bacterial pellets were harvested by centrifugation and resuspended in 50 mL of lysis buffer (20 mM Tris pH 7.5, 500 mM NaCl) to which 1 protease inhibitor tablet was added (Roche). Bacterial cells were lysed by sonication with 10 s pulses for 10 min and cell debris was removed by centrifugation at 25,000 x g for 40 min and cleared cell lysate was filtered through 0.45 μm filters. Filtered cell lysate was applied to 10 mL of Ni-NTA resin (Qiagen) which had been previously equilibrated with 10 column volumes of H2O, followed by 10 column volumes of lysis buffer. The resin was washed with 6 column volumes of lysis buffer followed by 6 column volumes of wash buffer (20mM Tris pH 7.5, 500 mM NaCl, 5 mM Imidazole). Full length TcdB toxins were eluted using 250 mM imidazole in 20mM Tris pH 7.5, 500 mM NaCl. The desired fractions were pooled and exchanged into 20 mM Tris pH 7.5, 40 mM NaCl and subjected to anion exchange chromatography using a 5 mL HiTrap Q anion exchange column (Cytiva). Proteins were fractionated using a gradient from 40 mM NaCl to 500 mM NaCl in 20 mM Tris pH 7.5 over 20 column volumes. Protein fractions were analyzed by SDS-PAGE and fractions containing full length TcdB variants were pooled and exchanged into 20 mM Tris pH 7.5, 150 mM NaCl and glycerol was added to a final concentration of 15% v/v. Protein was stored at -80 °C.

### TcdB-GTD glycosyltransferase assays

TcdB-GTD glycosyltransferase activity was measured using 6-^3^H UDP-glucose as a substrate and capturing radiolabeled glycosylated Rac1 protein by precipitating the protein product based on the method described in the study of Bensadoun and Weinstein^29^ and Paparella *et al* ^24,25^. Initial rate studies used fixed UDP-glucose/6-^3^H UDP-glucose or Rac1 while varying the concentration of the other substrate. Briefly, a glycosyltransferase reaction was carried out for at least 15 min at room temperature in a 50 μL reaction mix containing 50 mM HEPES pH 7.0, 100 mM, 4 mM MgCl_2_, 1 mM MnCl_2_ and purified TcdB-GTD (variants 1-8) at an appropriate concentration to detect the completed reaction in the linear phase (typically 1 nM, with the exception of TcdB4-GTD which was used at 90 nM). Terminated reaction samples were added to a precipitation mix of 120 μg/mL sodium deoxycholate, 6% trichloroacetic acid and 10 μg/mL BSA. Samples were incubated at room temperature for 15 min before being centrifuged for 20 min at 13,000 rpm to separate the glycosylated protein product. The supernatant was discarded and the protein pellet was resuspended in 500 μL of 200 mM Tris pH 7.5, 5% SDS and 20 mM NaOH. Samples were briefly vortexed and added to 20 mL liquid scintillation vials to which 10 mL of Ultima Gold was added (Perkin Elmer). The amount of glycosylated Rac1 was determined by liquid scintillation counting of each sample for 1 min using a Tri-carb 2910TR scintillation counter (Perkin Elmer). The initial rates of the reaction were fit to the Michaelis Menten equation (eq 1) to determine the Michaelis Menten constant (*K*_*m*_) for UDP-glucose and Rac1 for each TcdB-GTD variant.

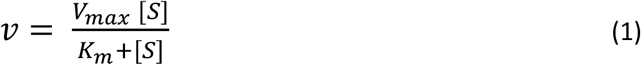

In kinetic inhibition assays, glycosylated Rac1 was analyzed as described above with the exception of 10 μM total UDP-glucose concentration (including 6-^3^H UDP-glucose), while the Rac1 was added at the *K*_*m*_ for each TcdB-GTD variant (Table 1). The IC_50_ value of UDP and isofagomine was determined from a dose-response curve by varying the concentration of the inhibitor under the same enzyme concentration. The data was analyzed with GraphPad Prism software using a non-linear fit of log10 [inhibitor] vs. normalized response. The inhibition constant (*K*_*i*_) for each compound was calculated using Eq. 2^31^, where S represents the total substrate concentration (10 μM) and *K*_*m*_ is the Michaelis-Menten constant for UDP-glucose for the respective TcdB-GTD variant (Table 1).

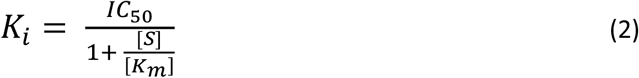

### Cell rounding assays

Cell rounding assays were performed using HeLa cells grown in DMEM medium with 10% FBS in 5% CO2 at 37 °C. Cells were acquired from the ATCC and maintained with standard trypsin passaging protocols. For quantitation of cell rounding, HeLa cells were seeded into 24-well plates at 40,000 cells/well. The following day, samples were treated with isofagomine (Serial dilution from 200 μM to 0.78 μM final concentration) and incubated for 15 minutes before addition of toxin. Cells were treated for 3-5 hours with 1 pM full length TcdB1/2, 10 nM TcdB3 and 40 pM TcdB4 at 37 °C, 5% CO2 before imaging. Samples were imaged by light microscopy on a Zeiss Axioscope with 4 representative images taken for each sample. Rounded cells were counted for each image and expressed as a percentage of total cells. Images were processed in FIJI/Image J, with manipulation limited to alterations of brightness and contrast. The data was analyzed with GraphPad Prism software using a non-linear fit of log10 [inhibitor] vs. normalized response to calculate the IC_50_ of isofagomine.

### Mouse toxigenic model

All procedures involving animals were conducted under protocols approved by the Institutional Animal Care and Use Committee at the Albert Einstein College of Medicine. The toxigenic model was based on the protocol described in Bender *et al*^26^. C57BL/6 mice (6 per group) at 10-12 weeks were provided drinking water that contained isofagomine tartrate (20 mg/kg, 60 mg/kg or 180 mg/kg). The control group was provided with drinking water without isofagomine tartrate (0 mg/kg). One day after being provided isofagomine, mice were intraperitoneally injected with TcdB1 (2 μg/kg in a volume of 100 μL). Mice continued to receive isofagomine tartrate *ad* libitum. Mice were monitored for up to 2 days post-challenge for toxic effect and were euthanized if they became moribund. Survival plots were created using GraphPad prism with death as the point when animals became moribund (time of sacrifice).

### Mouse *C. difficile* infection model

All procedures involving animals were conducted under protocols approved by the Institutional Animal Care and Use Committee at the Albert Einstein College of Medicine and the Royal Veterinary College (London). The *C. difficile* infection model was based on the mouse model described in Chen *et al*^27^. C57BL/6 mice (6 per group) at 13-14 weeks received antibiotic treatment in the drinking water for 3 days as shown in Figure 4. The antibiotic mixture contained: kanamycin (0.4 mg/mL), gentamicin (0.035 mg/mL), colistin (850 U/mL), metronidazole (0.215 mg/mL) and vancomycin (0.045 mg/mL). This corresponds to approximate daily doses of kanamycin (40 mg/kg), gentamicin (3.5 mg/kg), colistin (4.2 mg/kg), metronidazole (21.5 mg/kg) and vancomycin (4.5 mg/kg). After standard drinking water for 2 days following antibiotic treatment, mice were administered clindamycin (10 mg/kg) via oral gavage. On this same day, mice were also switched to drinking water containing isofagomine-tartrate (5 mg/kg, 15 mg/kg or 45 mg/kg) while the control group continued to receive standard drinking water. The following day, mice were orally administered 10^8^ CFU of *C. difficile* spores from the VPI 10463 strain (TcdA^+^ TcdB^+^, ribotype 087). Mouse weights and development of disease symptoms were monitored daily. Symptoms were monitored by visual examination 3-times per day and scored as follows: Grade 0 (hard fecal pellet, normal behavior), Grade 1 (difficulty in producing feces, mild weakness), Grade 2 (loose stool, weakness) and Grade 3 (liquid feces, inflamed rectum, wet (soiled) tail, extreme weakness). Mice that reached Grade 3 and/or lost >20% of their body weight were culled. Fecal samples were also collected at different time points during the experiment for microbiota analysis (see below). For histological analysis, colon tissue was removed and preserved in formalin for subsequent histopathology. For infected animals that were culled, colon tissue was removed from 2 animals randomly chosen per group. For surviving animals, at the study end, colon tissue was removed from 1 animal randomly chosen per group.

### Microbiota analysis

To analyze the GI microbiota, mouse fecal samples were collected at different time points during the CDI mouse model, described above. Fecal samples were collected at day -6 (5 samples taken randomly, 1 sample from each group), day -2 (before administration of isofagomine, 5 samples, 1 sample from each group) and on day 10 (Samples taken from each surviving mouse). Fecal samples (∼100mg) were processed using the Zymo Quick-DNA Fecal/Soil Microbe Miniprep Kit (D6010) following the manufacturer’s instructions and the purified DNA was sent to Azenta Life Sciences for sequencing.

## Author Contributions

A.S.P performed protein expression and purification, kinetic analysis, inhibition assays and cell rounding assays. A.S.P and I.B evaluated isofagomine using mouse intraperitoneal injection model. W.F, H.H and S.C evaluated isofagomine using *C. difficile* infection mouse model and coordinated fecal microbiome analysis. M.P, F.L and P.C.T performed synthesis of isofagomine. V.L.S designed and supervised the project. All authors contributed to writing the manuscript.

## Funding

This work was supported by NIH research grant and AI150971.

## Notes

### Competing Interest Statement

The authors have declared no competing interest.

