## Supplementary Figure 1 for "Isofagomine inhibits multiple TcdB variants and protects mice from *Clostridioides difficile* induced mortality"

### Supplementary Information

A. **Supplementary Methods**
B. **Supplementary Figures S1-S2**
C. **Supplementary Table 1**
D. **Supplementary References**

### A. Chemical Synthetic Schemes

#### Synthesis of isofagomine

Isofagomine was synthesized as reported (Rybczynski, P.; Tretyakov, A.; Fuerst, D.; Sheth, K. New method for preparing isofagomine and it's derivatives, US Patent 2010/0160638) with some modifications. Nitrile **1** (prepared from D-arabinose) was reduced with $BH_3.SMe_2$ to the amine which was isolated as it's N-Boc derivative **2**. This was then treated with aq HCl followed by hydrogenolysis affording isofagomine **3**.

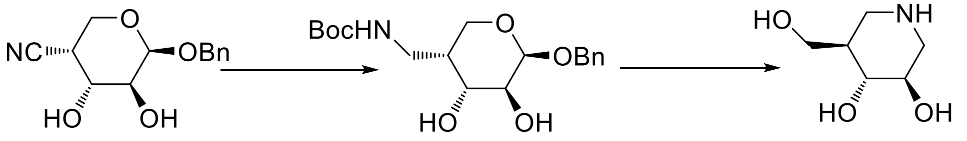


**1** **2** **3**

A solution of **1** (0.507 g, 2.03 mmol) in dry THF (10 mL) was stirred at RT under argon and borane dimethyl sulphide complex (1.03 mL, 0.822 g, 10.2 mmol) was added, then the solution was heated under reflux for 16 h. Methanol (8 mL) was carefully added and the solution was heated under reflux for 3 h and then concentrated to dryness. Methanol was added to the residue and the mixture concentrated to dryness three times. Then methanol (20 mL) was added followed by 10% aq sodium carbonate solution (5 mL). Di-tert-butyl dicarbonate 1.35 g, 6.10 mmol) was added and the mixture was stirred for 1 h and then concentrated to dryness. Chloroform (30 mL) was added and the mixture was filtered through celite and the solids washed with chloroform. Evaporation of the filtrates gave a solid. This was triturated with hexanes, filtered and dried to give **2** (0.62 g, 1.75 mmol, 86%) with NMR spectra consistent with that reported (Andersch, J.; Bols, M. *Chem. Eur. J.*, **2001**, *7*, 3744-3747).

To a solution of **2** (9.80 g, 27.7 mmol) in THF (100 mL) was added 5M aq HCl (75 mL) and the solution was stirred at RT for 2 h and then concentrated to dryness. A solution of the residue in methanol was treated with Amberlyst A26 (OH-) resin until the solution pH was > 7, and then filtered and concentrated to dryness. The residue in THF (100 mL) and methanol (100 mL) was stirred under hydrogen in the presence of 10% Pd/C for 16 h. After removal of the solids and solvent and chromatography of the residue DCM/MeOH/aq $NH_3$ 5:4:1) afforded isofagomine **3** (2.80 g, 19.0 mmol, 68%) which could be converted to the tartrate salt as reported. (Patent above).

#### Isothermal Titration Calorimetry

All ITC experiments were conducted on a Microcal PEAQ-ITC (Malvern Instruments). The experiments were performed at 25 °C in a cell containing 280 µL of reaction mixture (50 mM HEPES pH 7.5, 100 mM KCl, 4 mM $MgCl_2$, 1 mM $MnCl_2$) and 40 µM TcdB4-GTD. The ligand was titrated into the protein solution over 19 injections of 2 µL of 6 s with a 150 s equilibration period between injections. For binding measurements isofagomine, 40 µM TcdB4-GTD was incubated with 1 mM UDP in the buffer described above for 30 min in the sample cell prior to titrating 500 µM isofagomine. The resulting data were fit to a model of one distinct binding site. The first injection for each sample was excluded from data fitting. Titrations were run past the point of enzyme saturation to correct for heats of dilution.

**B. <u>Supplementary Figures</u>**

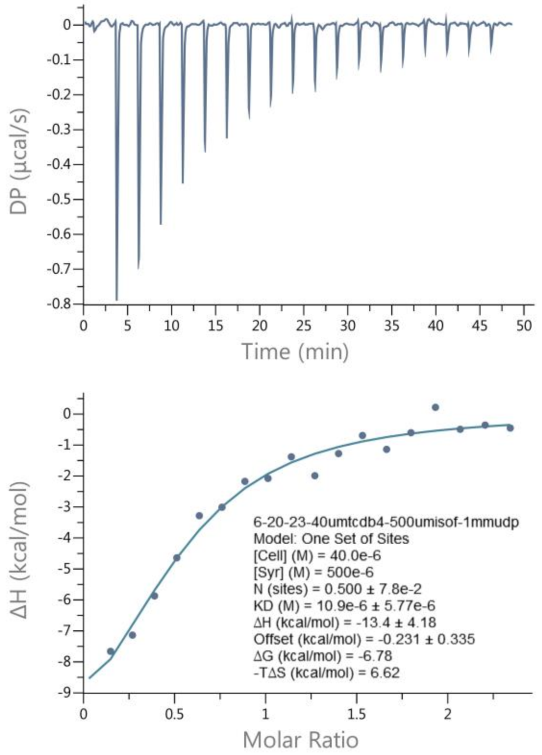


**Figure S1. ITC titration of Isofagomine binding to TcdB4-GTD in the presence of UDP.** Conditions are provided in supplementary methods. Top panel indicates raw heat data and bottom panel shows integrated heat injections which have been normalized per mole of injectant as a function of molar ratio. Data was fitted using a single binding site model.

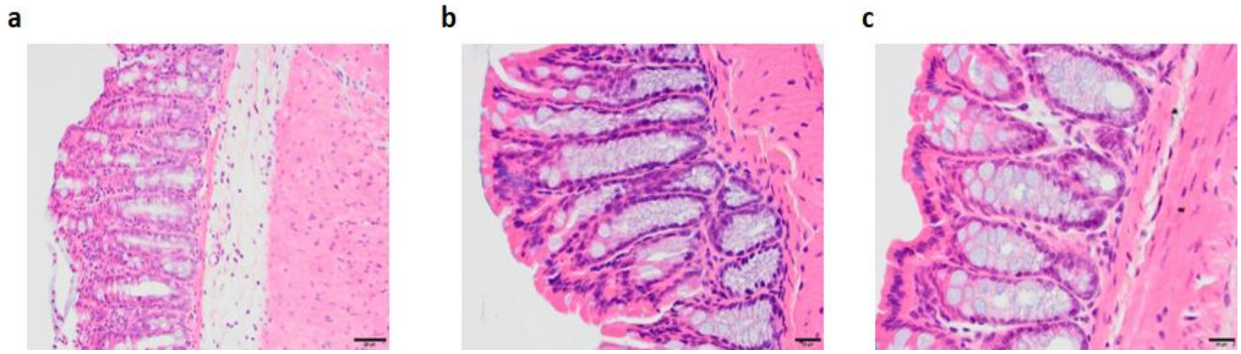


**Figure S2: Representative images (H&E stain) of colon tissue sections from treated and non-treated mice**. **a)** Non-treated mouse. **b)** Vancomycin treatment. **c)** 15 mg/kg isofagomine treatment. Black scale bars represent 50 µm.

### C. <u>Supplementary Table 1</u>

**Table S1: *Clostridioides difficile* strain information for TcdB variants used in this study**

| *C. difficile* Toxin | *C. difficile* strain name | Amino acid sequence length |
|---|---|---|
| TcdB1-GTD | 630 | 544 |
| TcdB2-GTD | CD196 | 544 |
| TcdB3-GTD | 1470 | 544 |
| TcdB4-GTD | 8864 | 544 |
| TcdB5-GTD | lsh26 | 544 |
| TcdB6-GTD | CD160 | 544 |
| TcdB7-GTD | CD10-165 | 544 |
| TcdB8-GTD | 173070 | 544 |
| TcdB1 Full length | 630 | 2366 |
| TcdB2 Full length | CD196 | 2366 |
| TcdB3 Full length | 1470 | 2367 |
| TcdB4 Full length | 8864 | 2367 |
